## Supporting Information for "Measuring context dependency in birdsong using artificial neural networks"

#### S1 Details on Syllable Clustering by VAE

This section is a detailed description of our syllable clustering based on ABCD-VAE (also see Morita and Koda, 2020, for another application of this proposed method). S1.1 describes the seq2seq backbone of the VAE, and S1.2 explains the ABCD-VAE together with the optimization objective. S1.3 defines the parameter settings and the training procedure. Finally, S1.4 discusses problems with the standard Gaussian VAE and the motivation behind our discrete VAE.

##### S1.1 Seq2Seq Autoencoder

Figure S1.1 shows the global architecture of our seq2seq VAE, which consists of three modules: the encoder, ABCD-VAE, and the decoder. The entire network receives a time series of syllable spectra, $\mathbf{y} := (\mathbf{y}_1, \dots, \mathbf{y}_T)$, as its input (in the encoder module) and reconstructs the input data. The reconstruction includes the prediction of each spectrum, $\hat{\mathbf{y}} := (\hat{\mathbf{y}}_1, \dots, \hat{\mathbf{y}}_T)$, as well as that of the offset $T$ of the time series, implemented by binary judgments of whether each time step $t$ is the offset ($h_t = 1$) or not ($h_t = 0$). During the reconstruction process, a fixed-dimensional representation of the entire syllable was obtained between the encoder and decoder and was classified into a discrete category by the ABCD-VAE module.

The backbone of the encoder module is the bidirectional LSTM (Hochreiter and Schmidhuber, 1997; Schuster and Paliwal, 1997). This RNN processes the input spectra forward and backward. The last hidden and cell states in the two directions are concatenated and transformed by a multi-layer perceptron (MLP).

The MLP output is fed to the ABCD-VAE module (see S1.2) and classified into a discrete category that has a corresponding real-value vector representation. The ABCD-VAE outputs the vector representation of the assigned category, which is concatenated with the embedding of the speaker $s$ of the input syllable. Accordingly, the discrete syllable categories in the ABCD-VAE need not encode speaker characteristics, resulting in speaker normalization (van den Oord et al., 2017; Chorowski et al., 2019; Dunbar et al., 2019; Tjandra et al., 2019). See S1.4 for the motivation behind using this speaker normalization for the birdsong data. The concatenation of the output from the ABCD-VAE and the speaker embedding is transformed by another MLP and fed to the decoder LSTM, which is unidirectional. For each time, step $t \in \{1, \dots, T\}$, the output from the LSTM is sent to two distinct MLPs. One of them computes the logits for the offset predictions ($\mathbb{P}(h_t)$). The other MLP parameterizes the isotropic Gaussian probability density function of the spectrum reconstruction (cf. Kingma and Welling, 2014). We sampled $\hat{\mathbf{y}}_t$ using this Gaussian, which is used as the input to the LSTM at the next time step $t + 1$ (the initial input is $\hat{\mathbf{y}}_0 = 0$).[1]

##### S1.2 ABCD-VAE

This section provides details on the ABCD-VAE. We start with a mathematical description of the model and then move to an explanation of the network implementation. Just like other VAEs, we assumed a prior distribution of the latent feature $z^{(i)}$ of each time-series data $i$. $z^{(i)}$ is discrete in this study and its prior is the Dirichlet-Categorical distribution. Eq. 1 and 2 below define this prior as the two-step generative procedure. The time-series data—represented by the spectra $\mathbf{y}^{(i)}$ and offset judgments $\mathbf{h}^{(i)}$ —are generated conditioned on $z^{(i)}$, whose probability function is implemented by the decoder (Eq. 3).

$$\boldsymbol{\pi} \sim \text{Dirichlet}(\boldsymbol{\alpha}) \tag{1}$$

$$z^{(i)} \mid \boldsymbol{\pi} \sim \text{Categorical}(\boldsymbol{\pi}) \tag{2}$$

$$(\mathbf{y}^{(i)}, \mathbf{h}^{(i)}) \mid z^{(i)} \sim p(\cdot \mid z^{(i)}, s^{(i)}) = \text{Decoder}(z^{(i)}, s^{(i)}) \tag{3}$$

[1]Note that the mathematically correct input to the decoder LSTM is the ground truth spectra, $\mathbf{y}_t$, rather than the reconstruction, $\hat{\mathbf{y}}_t$, because the objective function is the joint probability of $\mathbf{y}$ and $\mathbf{h}$ (see Eq. 4). However, the ground truth input to the decoder caused a *uninformative latent variable problem.* The decoder LSTM is considerably powerful and easily trained to fit to the overall distribution of the time-series data while ignoring information from the encoder (Bowman et al., 2016; Zhao et al., 2017; Liu et al., 2019). We found that our seq2seq VAE did not suffer from this issue when the noisy, reconstructed values $\hat{\mathbf{y}}_t$ were used instead of the ground truth (cf. also adopted by Chorowski et al., 2019).

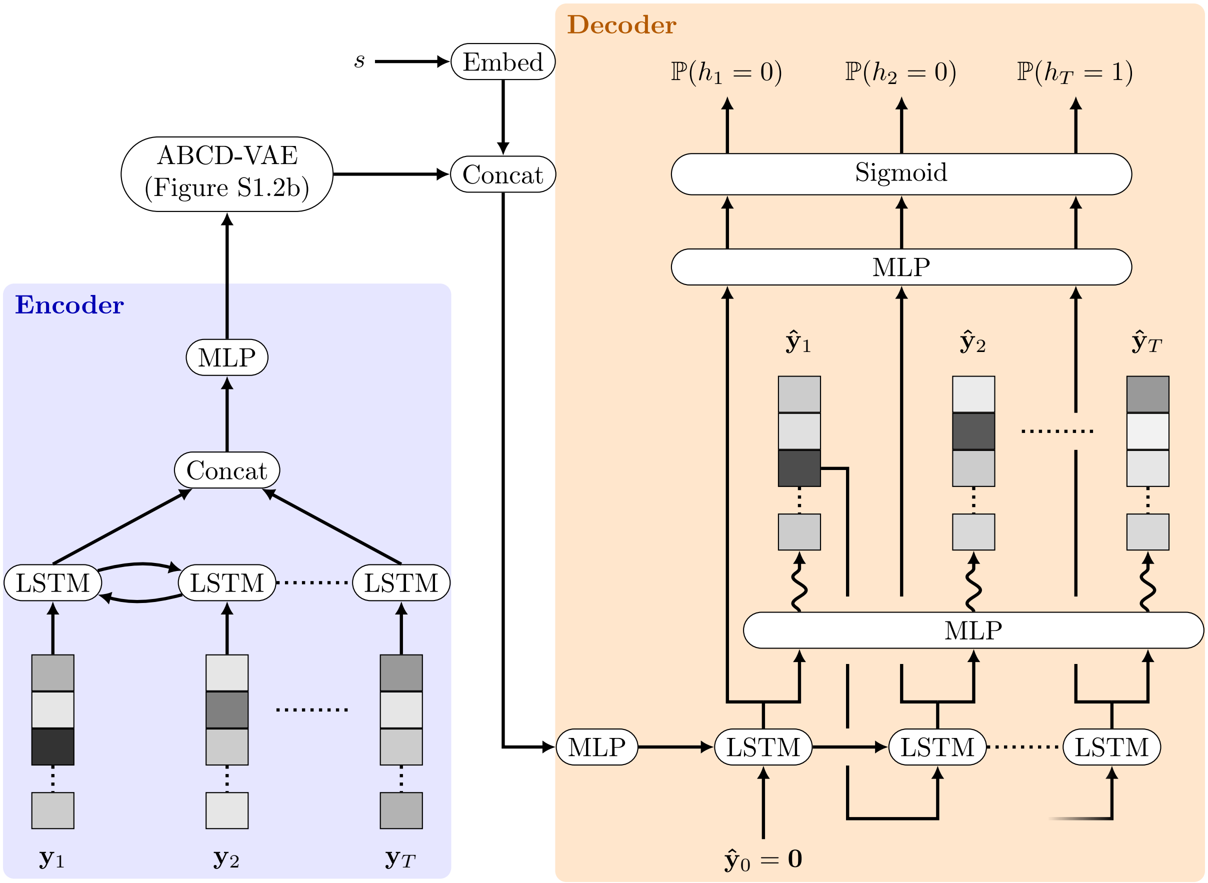


Figure S1.1: The architecture of the RNN-VAE, consisting of the Encoder module and the RNN Decoder module. The wavy arrows represent isotropic Gaussian sampling parameterized by the output of the previous MLP.

Where $\boldsymbol{\alpha} := (\alpha_1, \ldots, \alpha_K)$ are positive real numbers and $\boldsymbol{\pi} := (\pi_1, \ldots, \pi_K) \in \Delta^{K-1}$ is a probability vector (i.e., $\forall k \in \{1, \ldots, K\}, \pi_k \geq 1 \wedge \sum_{k=1}^{K} \pi_k = 1$). The Dirichlet-Categorical prior causes a rich-gets-richer bias, preferring a smaller number of categories to be used repeatedly (Bishop, 2006; O'Donnell, 2015; Little, 2019), while the (uniform) categorical prior—standard in the categorical VAE (Jang et al., 2017)—eats up all the categories available. Because of this Occam's razor effect, the Dirichlet-Categorical prior (and its extension to unbounded choices, the Dirichlet process) is popular in Bayesian learning when the model needs to detect the appropriate number of categories in the posterior (see Anderson, 1990; Kurihara and Sato, 2004, 2006; Teh et al., 2006; Kemp et al., 2007; Goldwater et al., 2009; Feldman et al., 2013; Kamper et al., 2017; Morita and O'Donnell, To appear, for examples in computational linguistics and cognitive science). The role of the encoder is to approximate the posterior $p(\boldsymbol{\pi}, \mathbf{z} \mid \mathbf{y}, \mathbf{h})$ of the model in Eq. 1-3. We make several assumptions on the approximated posterior, denoted by $q(\boldsymbol{\pi}, \mathbf{z} \mid \mathbf{y})$.[2]

1. $\boldsymbol{\pi}$ and $\mathbf{z}$ are independent given $\mathbf{y}$ in $q$: i.e., $q(\boldsymbol{\pi}, \mathbf{z} \mid \mathbf{y}) = q(\boldsymbol{\pi} \mid \mathbf{y}) q(\mathbf{z} \mid \mathbf{y})$.

2. Each $z^{(i)}$ is the categorical distribution and is independent of the other $z^{(j)}$ $(i \neq j)$ given the corresponding data $\mathbf{y}^{(i)}$.

3. $\boldsymbol{\pi}$ is independent of $\mathbf{y}$ in $q$: i.e., $q(\boldsymbol{\pi} \mid \mathbf{y}) = q(\boldsymbol{\pi})$.

4. $q(\boldsymbol{\pi})$ is the Dirichlet distribution whose parameters are in the form of $\boldsymbol{\omega} := N\boldsymbol{\theta} + \alpha$, where $N$ is the data size (the total number of time-series data) and $\boldsymbol{\theta}$ is a trainable vector in the simplex (i.e.,

[2] Note that the time series input into the encoder contains implicit information about its length/offset, $\mathbf{h}^{(h)}$.

$\forall k \in \{1, \dots, K\}, \theta_k \geq 0 \wedge \sum_k \theta_k = 1)$.

The assumptions in 1 and 3 are imported from the mean-field variational inference, and 3 provides the optimal form of $q(\boldsymbol{\pi})$ under the assumptions (Bishop, 2006).[3] The optimization objective of the entire VAE is the maximization of the evidential lower bound (ELBO) of the log marginal likelihood $\log p(\mathbf{y}, \mathbf{h})$ (Kingma and Welling, 2014; Bowman et al., 2016).

$$\begin{aligned}\log p(\mathbf{y}, \mathbf{h}) &\geq \log p(\mathbf{y}, \mathbf{h}) - D_{\mathrm{KL}}\left[q(\boldsymbol{\pi}, \mathbf{z} \mid \mathbf{y}) \parallel p(\boldsymbol{\pi}, \mathbf{z} \mid \mathbf{y}, \mathbf{h})\right] \\ &= - D_{\mathrm{KL}}\left[q(\boldsymbol{\pi}, \mathbf{z} \mid \mathbf{y}) \parallel p(\boldsymbol{\pi}, \mathbf{z})\right] + \mathrm{E}_q\left[\log p(\mathbf{y}, \mathbf{h} \mid \mathbf{z})\right] \\ &=: \mathrm{ELBO}\end{aligned} \tag{4}$$

Based on the assumptions of the approximated posterior $q$, the first term in Eq. 4 is rewritten as follows:

$$D_{\mathrm{KL}}\left[q(\boldsymbol{\pi}, \mathbf{z} \mid \mathbf{y}) \parallel p(\boldsymbol{\pi}, \mathbf{z})\right] = \mathrm{E}_q\left[\log q(\boldsymbol{\pi})\right] - \mathrm{E}_q\left[\log p(\boldsymbol{\pi})\right] + \sum_{i=1}^{N} \left(\mathrm{E}_q\left[\log q(z^{(i)} \mid \mathbf{y}^{(i)})\right] - \mathrm{E}_q\left[\log p(z^{(i)} \mid \boldsymbol{\pi})\right]\right)$$

Where each term has a closed form. During the mini-batch learning, the first two terms outside the summation operation, the index $i$ of which denotes data, are multiplied by $B/N$, where $B$ is the batch size. For the second term in Eq. 4, $\mathrm{E}_q\left[\log p(\mathbf{y}, \mathbf{h} \mid \mathbf{z})\right] = \sum_{i=1}^{N} \mathrm{E}_q\left[\log p(\mathbf{y}^{(i)}, \mathbf{h}^{(i)} \mid z^{(i)})\right]$, we approximate the computation of the expectation using the Monte Carlo method (Kingma and Welling, 2014). We adopted the Gumbel-Softmax approximation proposed by Jang et al. (2017) because the exact sampling from the categorical distribution $q(z^{(i)} \mid \mathbf{y}^{i})$ is incompatible with gradient-based training.

$$\mathrm{E}_q\left[\log p(\mathbf{y}^{(i)}, \mathbf{h}^{(i)} \mid z^{(i)})\right] \approx \log p(\mathbf{y}^{(i)}, \mathbf{h}^{(i)} \mid \tilde{\mathbf{z}}^{(i)}) \qquad (\tilde{\mathbf{z}}^{(i)} \in \Delta^{K-1} : \text{Sample from Gumbel-Softmax})$$

To simplify the learning process, the linear transformation before and after the Gumbel-Softmax sampling share the same weight matrix $\mathbf{M}$. The linear transformation before the Gumbel-Softmax computes the logits by multiplying the output of the encoder with $\mathbf{M}$, while the transformation after the Gumbel-Softmax computes $\tilde{\mathbf{z}}^{(i)}\mathbf{M}^{\mathrm{T}}$ (Figure S1.2a).[4] In other words, $\mathbf{M}$ is the "codebook" whose column vectors are the real-value representation of the corresponding discrete categories. The first linear transformation computes the similarity between the encoder output and each column vector, and the second transformation picks up the column vector of the sampled category (assuming that the Gumbel-Softmax sample $\tilde{\mathbf{z}}^{(i)}$ is close to a one-hot vector). Thus, our VAE is similar to the vector-quantized VAE (van den Oord et al., 2017; Chorowski et al., 2019; Tjandra et al., 2019), which uses the L2-similarity instead of our unnormalized cosine similarity (without the randomness).

The exact implementation of the ABCD-VAE is based on the scaled dot-product attention used in the Transformer (single head attention; Vaswani et al., 2017; Devlin et al., 2018) and depicted in Figure S1.2b. The attention mechanism first computes the dot product of the encoder's output (query) and the codebook $\mathbf{M}$ (memory; i.e., both key and value), yielding the similarity between the two. This similarity is scaled by $\sqrt{D_h}$, where $D_h$ is the dimensionality of the encoder output and the column vectors of the codebook (i.e., the number of rows in $\mathbf{M}$). The scaled similarity is transformed by the softmax into the posterior probability vector $q(z^{(i)} \mid \mathbf{y}^{(i)})$, and a Gumbel-Softmax sample $\tilde{\mathbf{z}}^{(i)}$ is drawn from this probability distribution. Finally, $\tilde{\mathbf{z}}^{(i)}$ is multiplied by $\mathbf{M}^{\mathrm{T}}$ (used as the value) and sent to the decoder.

#### S1.3 Parameter Settings and Training Procedure

The input time series $\mathbf{y}_1, \dots, \mathbf{T}$ were obtained as follows. We first applied the short-time Fourier transform to the recordings having the 8 msec Hanning window and 4 msec stride (i.e., 256 samples for the window

[3]The optimal update of $\boldsymbol{\theta}$ is proportional to the expected number of data classified into each category (Bishop, 2006). However, the exact computation of these expected counts requires iterations over all data and is inefficient. Therefore, we train $\boldsymbol{\theta}$ using the gradient ascent.

[4]For simplicity, the linear transformations do not have bias terms.

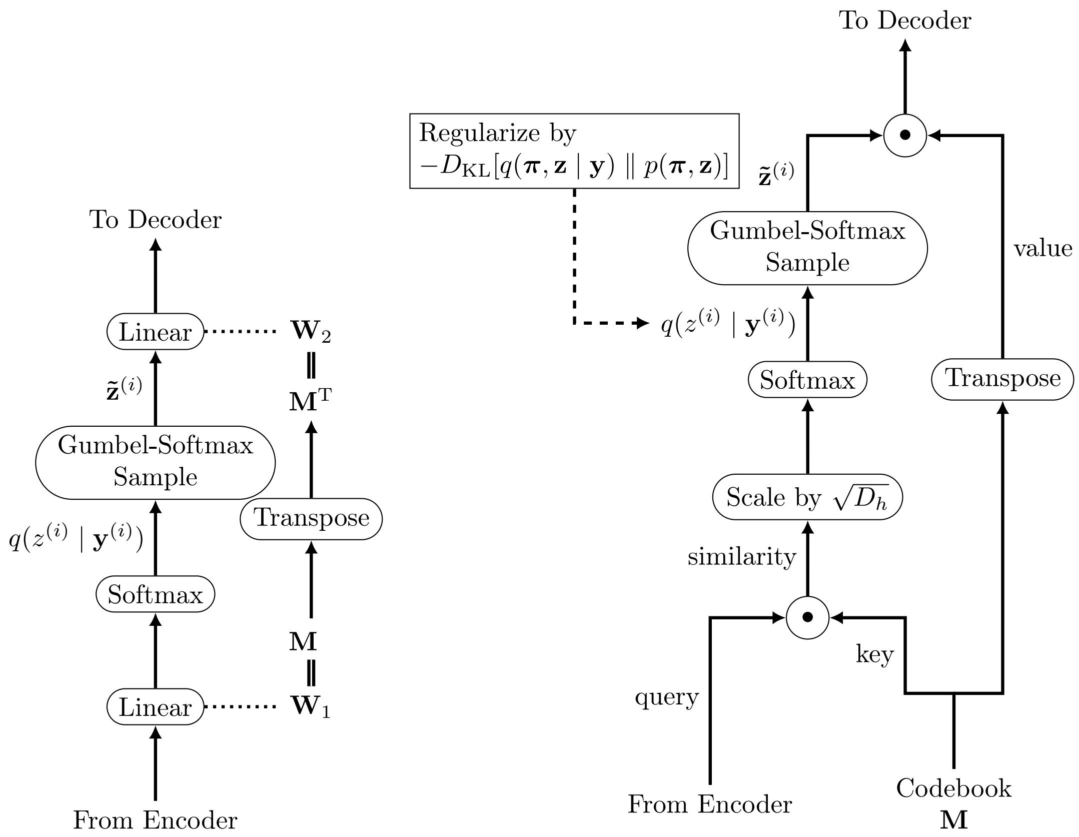


(a) Shared Weight Matrix

(b) Attention-based Implementation

Figure S1.2: ABCD-VAE.

length and 128 samples for the step size because the sampling rate was 32 kHz). The spectral amplitude was then log-transformed (after $2^{-15}$ was added to avoid underflow) and rescaled by $11^{-1}$ ($> \log 2^{15}$).

All the hidden states in the VAE had the dimensionality $D_h = 256$. The number of possible discrete categories (i.e., upper bound) was $K = 128$ and the non-linearity of the MLPs was tanh. The neural network was trained by the stochastic gradient ascent for 20 epochs. The first five epochs were used for "pretraining" where the posterior probability $q(z^{(i)} \mid \mathbf{y}^{(i)})$ was multiplied with the codebook $\mathbf{M}$ without sampling from the Gumbel-Softmax distribution. After this process, the temperature $\tau$ of the Gumbel-Softmax was annealed every 1000 iterations according to the schedule $\tau = \exp(-10^{-5}m)$, where $m$ is the number of iterations after the initial five epochs (cf. Jang et al., 2017). The learning rate was initially set as 1.0 and multiplied by 0.1 after every epoch in which validation loss was not improved. The gradient norms were clipped at 1.0 to avoid an explosion. No dropout or momentum were introduced.

##### S1.4 Problems with the Gaussian VAE Analysis of Birdsong

This section discusses the real-value features and problems of birdsong syllables obtained by the standard Gaussian VAE. The Gaussian VAE is a popular way of obtaining real-valued features of data in an arbitrary dimensional space (Kingma and Welling, 2014) and has been used for analyses of animal vocalization (Coffey et al., 2019; Goffinet et al., 2019; Sainburg et al., 2019b). We also tested it on our birdsong data and reported clustering results based on the acoustic features extracted from it as baselines (see Table 1 in the main text). The syllable features of the Gaussian VAE exhibited some clear clusters when we looked at each individual bird separately (Figure S1.3a, the original features had 16 dimensions, which we embedded into the two-dimensional space by tSNE for visualization, van der Maaten and Hinton, 2008).

When we represent syllables of multiple individuals, however, we no longer see such clear clusters. Figure S1.3b shows the syllable features of all the 18 Bengalese finches used in this study (distinguished by the colors and shapes of the markers). Many syllables are confused around the center, and tiny clusters in the peripheral are individual-specific.

We also tried the speaker normalization technique used in discrete VAEs (van den Oord et al., 2017; Chorowski et al., 2019; Tjandra et al., 2019), feeding the speaker ID the decoder module (Figure S1.1, wherein the speaker embeddings were jointly trained with the other modules; cf. Louizos et al., 2016), but this did not solve the problem (Figure S1.3c); GMM clustering in the resulting feature space aligned poorly with human annotations when the number of syllable categories was manually specified as 14 (= the annotation labels) and 37 (= the ABCD-VAE result), and resorted to the fine-grained, speaker-specific classification when the number of categories was auto-detected ($\geq 128$; Table S1.1). The speaker normalization increased the speaker perplexity only when the number of syllable categories was fixed as moderate numbers. That is, the speaker normalization only removed the global speaker-based zoning of the latent space, and individuality was still visible locally to the extent that the uncertainty of speaker identification from a 128-wise classification was less than a flip of a coin. Given the inappropriate distribution of the syllable features encoded by the Gaussian VAE (but see Louizos et al., 2016; Ganin et al., 2016, for other individual-normalization techniques that work on continuous spaces), we adopted end-to-end clustering with the ABCD-VAE.

##### S1.5 V-measure Metric for Evaluation of Clustering Results

In the main text, we evaluated the clustering results of birdsong syllables by their alignment with human annotations measured by Cohen's Kappa coefficients and homogeneity. These two metrics examined whether clustered syllables were annotated with the same label. On the other hand, they did not penalize overclassification. In our study, we compared the baseline and topline results with different numbers of syllable categories, so our evaluation did not take into account overclassification. In more general situations, however, overclassification is an important aspect of clustering and thus must be penalized. One option for a more comprehensive metric of clustering quality is *V-measure*, which combines homogeneity with another submetric for scoring overclassification, called *completeness*. Completeness evaluates clustering results in the opposite direction from homogeneity: it requires syllables annotated with the same label to belong to the same model-predicted category. For example, suppose that syllables labeled as "a" are split into two categories, "A" and "B", by clustering. As explained in the main text, homogeneity is satisfied even under this overclassification, as long as all the syllables in "A" and "B" are labeled as "a" and nothing else. By contrast, the classification violates completeness and, thus, it complements homogeneity as an evaluation

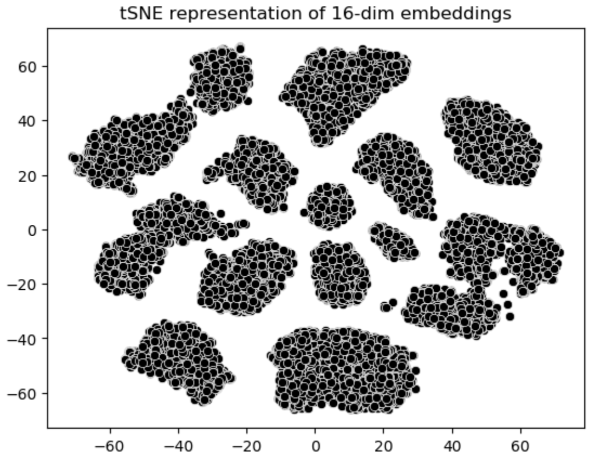


(a) Single Individual (b03)

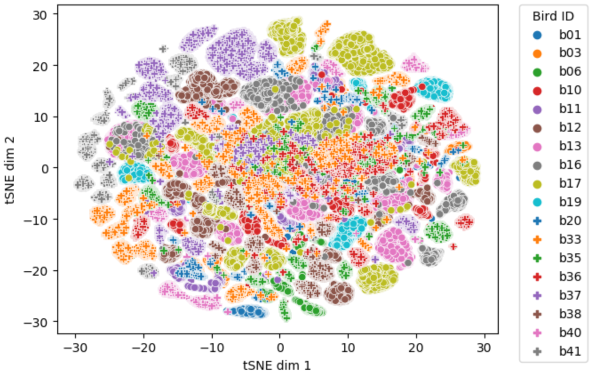


(b) All Individuals

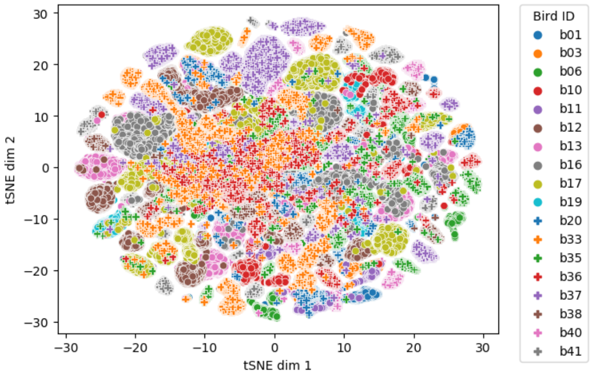


(c) Speaker-Normalized

Figure S1.3: Continuous-valued encodings of Bengalese finch syllables by Gaussian VAE. The original dimensionality of the features was 16, and they were embedded into the two-dimensional space by tSNE. (a) Syllables from a single individual (b03). (b) Syllables from all the individuals were plotted together, using colors and shapes of the markers to distinguish the individuals. (c) Syllables from all the individuals encoded with the speaker normalization technique used in discrete VAEs.

Table S1.1: Quantitative evaluation of the clustering by the ABCD- vs. Gauss-VAE for Bengalese finch syllables, extending Table 1 with the results based on the speaker-normalized Gauss-VAE. Cohen's kappa coefficient and homogeneity evaluated the alignment of the discovered clusters with manual annotations by a human expert. These scores for each individual bird were computed separately and their mean, maximum, and minimum over the individuals were reported since the manual annotation was not shared across individuals (see Method). Additionally, the perplexity of individual identification scored the amount of individuality included in the syllable categories yielded by the ABCD- and Gauss-VAE. The best scores are in boldface (results under the all-birds-together and bird-specific settings were ranked separately).

| Data | Method | # of clusters (source) | Cohen's Kappa mean [min,max] | Homogeneity mean [min,max] | Speaker Perplexity |
|---|---|---|---|---|---|
| All-Birds-Together | ABCD-VAE | 37 | **0.8990** [**0.7740**, **0.9929**] | **0.9084** [**0.7635**, 0.9868] | 8.0434 |
| All-Birds-Together | Gauss-VAE + GMM | 37 (ABCD-VAE) | 0.7446 [0.5956, 0.8912] | 0.7844 [0.6004, 0.9086] | 4.0783 |
| All-Birds-Together | Gauss-VAE + GMM | 14 (manual) | 0.6057 [0.4250, 0.8972] | 0.6718 [0.5053, 0.8536] | 6.7212 |
| All-Birds-Together | Gauss-VAE + GMM | ≥128 (auto-detected) | 0.8475 [0.5725, 0.9911] | 0.8773 [0.6666, 0.9869] | 1.7112 |
| All-Birds-Together | Gauss-VAE + GMM + Speaker-Normalization | 37 (ABCD-VAE) | 0.6403 [0.4410, 0.8471] | 0.6529 [0.4714, 0.8367] | 9.0148 |
| All-Birds-Together | Gauss-VAE + GMM + Speaker-Normalization | 14 (manual) | 0.4820 [0.2787, 0.6497] | 0.4975 [0.2863, 0.6600] | **11.3122** |
| All-Birds-Together | Gauss-VAE + GMM + Speaker-Normalization | ≥128 (auto-detected) | 0.8649 [0.5320, 0.9885] | 0.8754 [0.5800, **0.9904**] | 1.9887 |
| Bird-Specific | Gauss-VAE + GMM | 37 (ABCD-VAE) | 0.9304 [0.6619, 0.9906] | 0.9292 [0.6479, 0.9893] | — |
| Bird-Specific | Gauss-VAE + GMM | 5–14 (manual) | 0.7888 [0.5012, 0.9328] | 0.8090 [0.4732, 0.9254] | — |
| Bird-Specific | Gauss-VAE + GMM | 50–109 (auto-detected) | 0.9516 [0.7629, 0.9982] | 0.9505 [0.7687, 0.9962] | — |
| Bird-Specific | Gauss-VAE + GMM + Speaker-Normalization | 37 (ABCD-VAE) | 0.9137 [0.6052, 0.9893] | 0.9152 [0.5952, 0.9893] | — |
| Bird-Specific | Gauss-VAE + GMM + Speaker-Normalization | 5–14 (manual) | 0.7641 [0.3636, 0.9271] | 0.7889 [0.4271, 0.9401] | — |
| Bird-Specific | Gauss-VAE + GMM + Speaker-Normalization | 61–104 (auto-detected) | **0.9560** [**0.8046**, **0.9943**] | **0.9557** [**0.8011**, 0.9949] | — |

metric. (Note that completeness does not penalize non-uniformity of clustered syllables regarding human annotations, which violates homogeneity; e.g., it is satisfied even when the model-predicted category "A" includes syllables annotated with different labels, say "a" and "b", as long as all the "a" and "b" syllables belong to "A" and nowhere else.) Mathematically, violation of completeness is defined by the conditional entropy of the distribution of predicted clusters $\mathcal{K}$ given the ground truth classes $\mathcal{C}$:

$$\text{completeness}(\mathcal{C}, \mathcal{K}) := \begin{cases} 1 & H(\mathcal{K}) = 1 \\ 1 - \frac{H(\mathcal{K} \mid \mathcal{C})}{H(\mathcal{K})} & \text{Otherwise} \end{cases}$$

Note that the non-conditional entropy $H(\mathcal{K})$ is the normalizing term that makes completeness range between 0 and 1. Finally, V-measure is defined by the harmonic mean of homogeneity and completeness:

$$\text{V-measure}(\mathcal{C}, \mathcal{K}) := \frac{2 * \text{homogeneity}(\mathcal{C}, \mathcal{K}) * \text{completeness}(\mathcal{C}, \mathcal{K})}{\text{homogeneity}(\mathcal{C}, \mathcal{K}) + \text{completeness}(\mathcal{C}, \mathcal{K})}$$

Table S1.2 reports the completeness and V-measure scores of the syllable clustering results. We also repeat homogeneity in Table 1 for easier comparison among scores.

##### S1.6 Classification Confidence against Variations in Duration and Amplitude

This section discusses the robustness of clustering by the ABCD-VAE regarding variations in the amplitude and duration of birdsong syllables. Figure S1.4–S1.7 plots the probability of MAP syllable categories of Bengalese finch (S1.4, S1.5) and zebra finch (S1.6, S1.7) against the maximum amplitude (S1.4, S1.6) and duration (S1.5, S1.7) of the syllables. We can see no general pattern such as "classification probability decreases (i.e., the model gets less confident) as the amplitude/duration diverges from the mean". Instead, different categories use amplitude/duration information differently, some exhibiting a tendency to assign a greater probability to shorter/longer or louder/softer syllables while the others are more robust against variations in amplitude/duration. Overall, the Pearson correlation coefficient between the MAP classification probability and the deviation of the amplitude/duration from the median (all in log scale) was small: -0.0791 (vs. amplitude) and 0.0559 (vs. duration) for Bengalese finch; 0.1217 (vs. amplitude) and 0.0328 (vs. duration) for zebra finch. It also is certain that none of the detected syllable categories was purely characterized by amplitude nor duration; syllables having the same amplitude/duration were assigned various classification probabilities, indicating the existence of other acoustic factors that have an effect on the classification.

We conclude this section by introducing a possible method to ignore variations in amplitude/duration as noise. Remember that the learning objective of the canonical VAE is to reconstruct each input syllable as precisely as possible, and we adopted this framework in the present study. However, one can also add some noise to the input—including manipulation of amplitude or duration (Moulines and Charpentier, 1990)—and train a *denoising* VAE to recover the original data (Vincent et al., 2008). Then, the VAE will ignore variations in the manipulated acoustic dimension as noise rather than the systematic differences among categories.

#### S2 Details on the Transformer Language Model

Our analysis of context dependency was based on the Transformer language model of a Bengalese finch song and English sentences (Vaswani et al., 2017; Devlin et al., 2018; Dai et al., 2019). This section discusses the model parameters and training procedure we used. The Transformer consisted of six layers with eight attention heads per layer. The dimensionality of the hidden states, including the middle layer of the MLPs, was 512. We adopted the relative position encoding proposed by Dai et al. (2019). The input embeddings of the Bengalese finch syllables were additively combined with the embeddings of the speaker identity. Dropout was applied at the rate of 0.1 to the input embeddings (+ the speaker embeddings for the Bengalese finch data), the output of each Transformer sublayer before the residual connection, and to the attention weights. We trained the Transformer for $20,000$ iterations using the Adam optimizer with a learning rate of 0.001, $\beta_1 = 0.9$, $\beta_2 = 0.999$, and weight decay of 0.01 (Kingma and Ba, 2015). The learning rate was updated according to the schedule used by Vaswani et al. (2017) and Devlin et al. (2018), with $1,000$ warmup iterations. The batch size was 128.

Table S1.2: Scores of the clustering by the ABCD-VAE. Homogeneity, completeness, and V-measure evaluated the alignment of the discovered clusters with manual annotations by a human expert. These scores for each individual bird were computed separately and their mean, maximum, and minimum over the individuals were reported since the manual annotation was not shared across individuals (see Method). The best scores are in boldface (results under the all-birds-together and bird-specific settings were ranked separately).

| Species | Method | # of clusters (source) | Homogeneity mean [min,max] | Completeness mean [min,max] | V-measure mean [min,max] |
|---|---|---|---|---|---|
| BF | ABCD-VAE | 37 | **0.9084** [**0.7635**, 0.9868] | 0.6859 [0.5069, 0.8468] | 0.7765 [**0.6575**, 0.8797] |
| | Gauss-VAE + GMM (All-Birds-Together) | 37 (ABCD-VAE) | 0.7844 [0.6004, 0.9086] | 0.7595 [0.5555, 0.9190] | 0.7679 [0.6105, **0.8940**] |
| | | 14 (manual) | 0.6718 [0.5053, 0.8536] | **0.8249** [**0.6503**, **0.9492**] | 0.7353 [0.6158, 0.8489] |
| | | ≥128 (auto-detected) | 0.8773 [0.6666, **0.9869**] | 0.7177 [0.5499, 0.8652] | **0.7837** [0.6432, 0.8743] |
| | Gauss-VAE + GMM (Bird-Specific) | 37 (ABCD-VAE) | 0.9292 [0.6479, 0.9893] | 0.5890 [0.4058, 0.7522] | 0.7148 [0.5750, 0.8494] |
| | | 5–14 (manual) | 0.8090 [0.4732, 0.9254] | **0.8038** [**0.5771**, **0.9216**] | **0.8042** [0.5255, **0.9235**] |
| | | 50–109 (auto-detected) | **0.9505** [**0.7687**, **0.9962**] | 0.6076 [0.4284, 0.7712] | 0.7357 [**0.5965**, 0.8647] |
| ZF | ABCD-VAE | 17 | 0.6793 [0.4972, 0.8718] | 0.6141 [0.3346, 0.8495] | 0.6378 [0.4189, 0.8605] |
| | Gauss-VAE + GMM (All-Birds-Together) | 17 (ABCD-VAE) | 0.6177 [0.3030, 0.8942] | 0.7351 [0.4159, 0.9461] | 0.6502 [0.3638, 0.8279] |
| | | 13 (manual) | 0.6315 [0.0433, 0.9609] | **0.7569** [0.4104, 0.9494] | 0.6687 [0.0793, 0.9330] |
| | | ≥128 (auto-detected) | **0.9016** [**0.7643**, **0.9894**] | 0.7542 [**0.4514**, **0.9935**] | **0.8118** [**0.6173**, **0.9511**] |
| | Gauss-VAE + GMM (Bird-Specific) | 17 (ABCD-VAE) | 0.9545 [0.8828, 0.9905] | 0.5792 [0.2983, 0.7835] | 0.7132 [0.4566, 0.8453] |
| | | 4–13 (manual) | 0.8623 [0.7056, 0.9607] | **0.7911** [**0.4526**, **0.9886**] | **0.8219** [**0.5858**, **0.9226**] |
| | | 18–47 (auto-detected) | **0.9782** [**0.9274**, **1.0000**] | 0.5587 [0.3172, 0.7408] | 0.7039 [0.4781, 0.8359] |

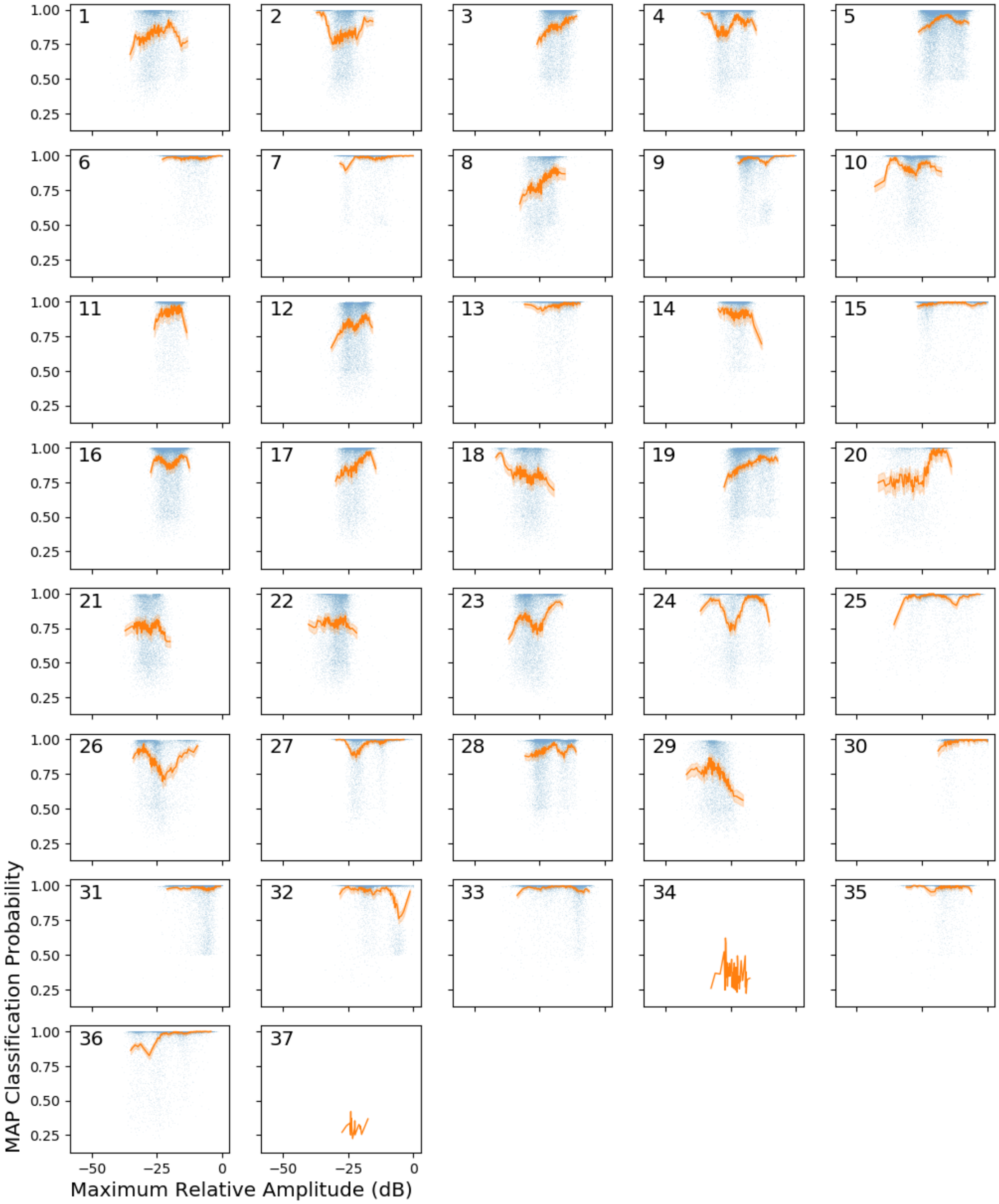


Figure S1.4: Probability of MAP syllable categories (indexed by the integer in the upper right) in Bengalese finch song against maximum amplitude of the syllables. The blue scatter represents individual syllables. The orange lines represent the mean probability over 1%tile windows.

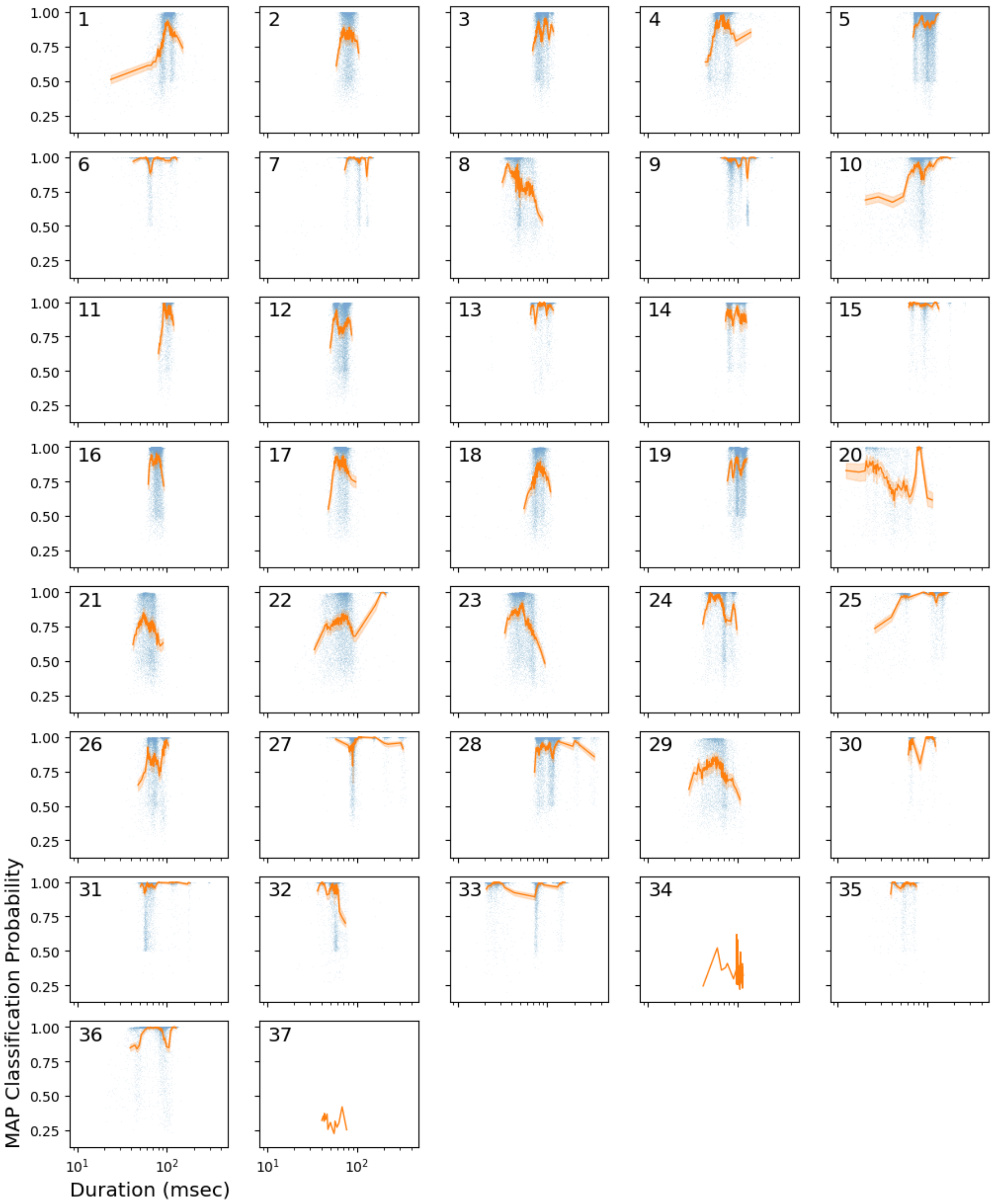


Figure S1.5: Probability of MAP syllable categories (indexed by the integer in the upper right) in Bengalese finch song against syllable duration. The blue scatter represents individual syllables. The orange lines represent the mean probability over 1%tile windows.

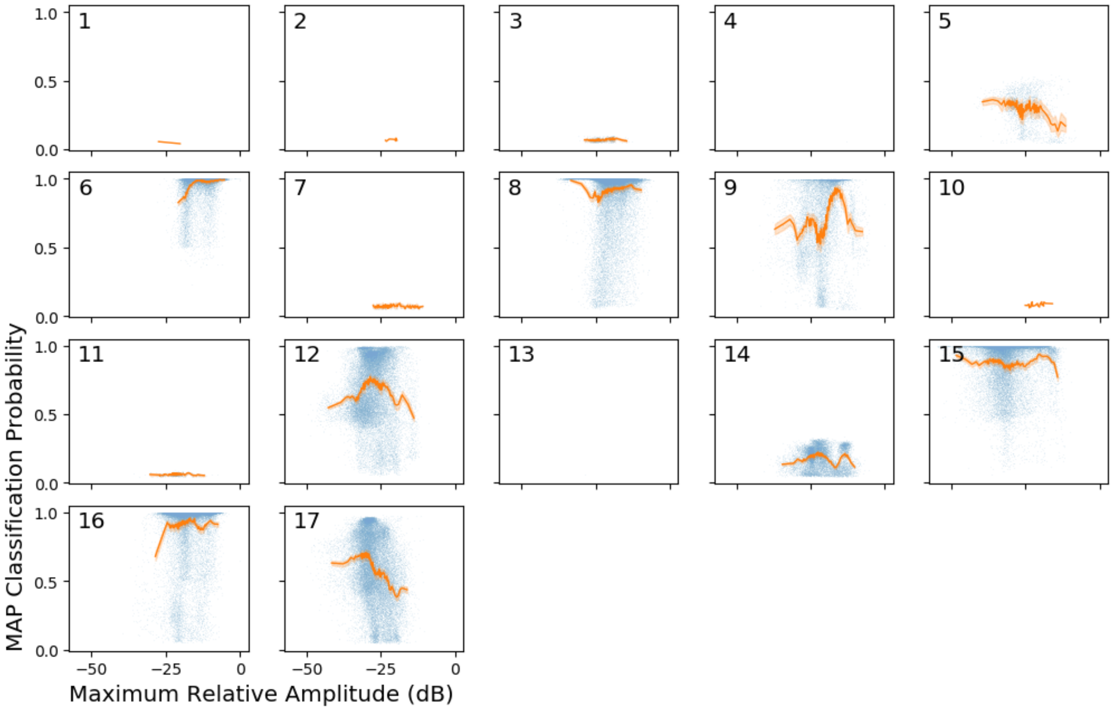


Figure S1.6: Probability of MAP syllable categories (indexed by the integer in the upper right) in zebra finch song against maximum amplitude of the syllables. The blue scatter represents individual syllables. The orange lines represent the mean probability over 1%tile windows. Note that categories numbered 4 and 13 were singletons and thus left empty.

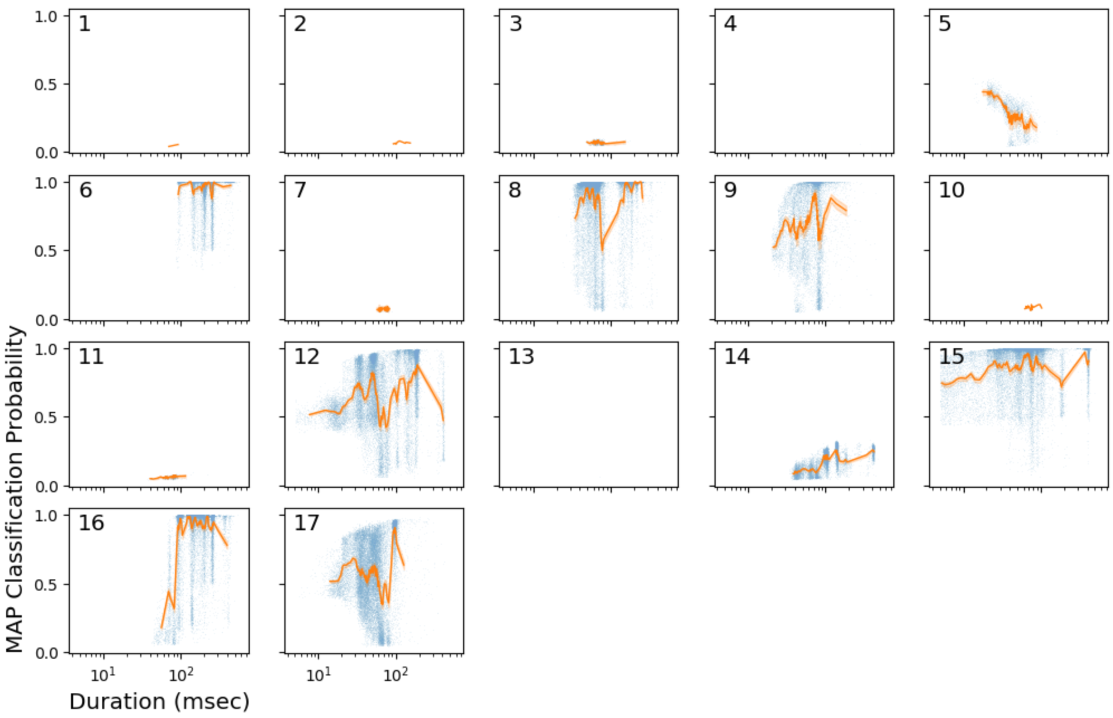


Figure S1.7: Probability of MAP syllable categories (indexed by the integer in the upper right) in zebra finch song against syllable duration. The blue scatter represents individual syllables. The orange lines represent the mean probability over 1%tile windows. Note that categories numbered 4 and 13 were singletons and thus left empty.

Table S3.1: The size of the training and test data used in the neural language modeling of zebra finch songs. The "SECL" portion of the test syllables was used to estimate the SECL. The numbers of syllables in parentheses report the incomplete syllables that were broken off at the start/end of recordings, which were labeled with a distinct symbol.

| Usage | # of sequences | # of syllables | |
|---|---|---|---|
| | | Total | SECL |
| Training<br>(incomplete) | 11,722 | 234,674<br>(5,763) | — |
| Test<br>(incomplete) | 100 | 2,936<br>(55) | 1,536<br>(49) |

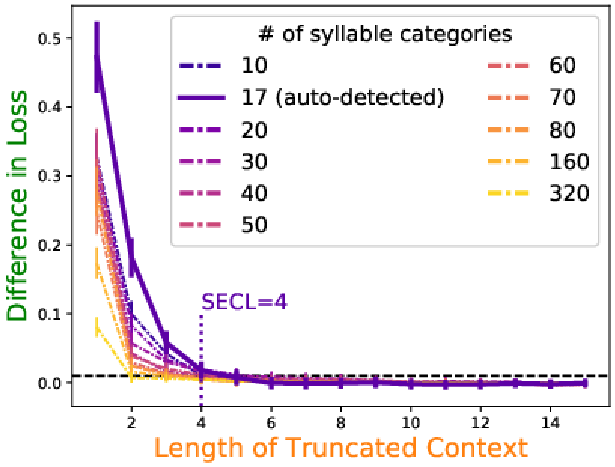


Figure S3.1: The differences in the mean loss (negative log probability) between the truncated- and full-context predictions of zebra finch songs. The x-axis corresponds to the length of the truncated context. The error bars show the 90% confidence intervals estimated from 10,000 bootstrapped samples. The loss difference is statistically significant if the lower side of the intervals are above the threshold indicated by the horizontal dashed line.

#### S3 Analysis of Context Dependency in Zebra Finch Song

This section reports the Transformer-based analysis of context dependency in zebra finch songs. Readers should remember that the unsupervised, speaker-invariant classification of zebra finches' syllables was not as reliable as Bengalese finches' and that the context dependency reported here is dependent on those classification results.

We used the same song data as those used in the unsupervised syllable clustering by the ABCD-VAE. The data consisted of 11,822 sequences of zebra finch syllables (each containing 1–219 syllables, 20.10 syllables on average) and 11,722 of them were used for training the Transformer language model (Table S3.1). The remaining 100 sequences were used to score the predictive performance of the trained model, from which the dependency (SECL) was calculated.

Our analysis estimated the SECL of zebra finch songs as four (Figure S3.1). Just as Bengalese finch songs did, zebra finch songs showed a trade-off between the number of syllable categories and context dependency, except that the seventeen-way classification—which was automatically detected by the ABCD-VAE—showed a greater difference than the ten-way classification. The difference between the model predictions based on the truncated and full contexts became smaller as the number of syllable categories increased (Fig.; $p < 0.001$ according to the linear regression of the loss difference on the number of syllable categories and the length of truncated contexts, both in the log scale).

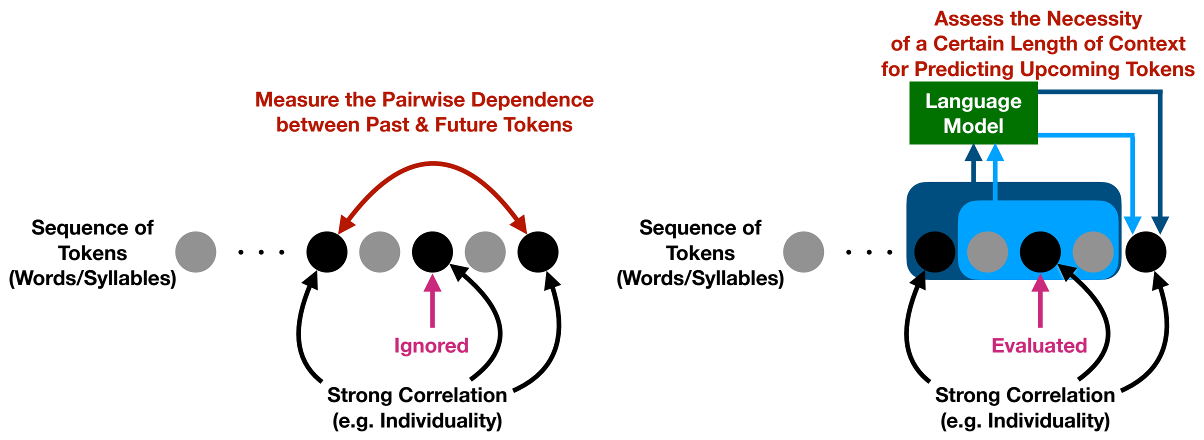


(a) (Naive) Mutual Information Approach

(b) Language Modeling Approach

Figure S4.1: The analysis of context dependency based on the (a) naive mutual information and (b) language modeling.

#### S4 Detailed Comparison with the Mutual Information Analysis

This section is a discussion of concrete examples of sequential data where the mutual information metric diverges from the intuitive concept of "context dependency," defined by or related to the memory burden on animal agents that produce/recognize the sequential data. We first introduce a naive definition of mutual information and show that individual-specific tokens can make it constant regardless of their distance in time series. We then discuss a modified version of mutual information that was adopted in previous studies of empirical data, providing some simulation results that demonstrate differences between the mutual information and model-based analyses regarding the detected context dependency (S4.2). It should be noted that recent studies on human language and birdsong did not use mutual information to assess the agent-based context dependency. Instead, they analyzed the decay in the mutual information to diagnose the generative model behind the data (Lin and Tegmark, 2017; Sainburg et al., 2019a).

##### S4.1 Problem with Individuality

The mutual information $I$ measures the expected divergence between the joint distribution of two tokens, $X$ and $X_{+d}$, at certain distance $d$ and the product of their marginal probability.

$$\begin{aligned} I(X, X_{+d}) :=& \mathrm{E}\left[\log_2 \frac{\mathbb{P}(X, X_{+d})}{\mathbb{P}(X)\mathbb{P}(X_{+d})}\right] \\ =& \sum_{x}\sum_{x_{+d}} \mathbb{P}(X = x, X_{+d} = x_{+d}) \log_2 \frac{\mathbb{P}(X = x, X_{+d} = x_{+d})}{\mathbb{P}(X = x)\mathbb{P}(X_{+d} = x_{+d})} \end{aligned} \tag{5}$$

Mutual information is zero if $X$ and $X_{+d}$ are independent. It should be noted that this is a pairwise metric, and the other tokens appearing between the two are ignored. For example, $X_{+d}$ can be *conditionally* independent of $X$ given other tokens between them, providing the same information as $X_{+d}$ for the prediction of $X$, but such relations are not detected by the metric (see Figure S4.1a).[5]

The mutual information, when naively defined as Eq. 5, diverges from the natural concept of context dependency when some tokens encode individual information. Suppose that sequences of tokens are generated by iterating the following procedure:

[5] It is not impossible to assess the conditional effect of mutual information in principle. We may replace all the probabilities in Eq. 5 with the conditional ones. Such an index, however, would be difficult to estimate in practice because of the exponentially possible sequences of conditioning tokens.

| Predecessor ($X$) | Follower ($X_{+d}$) | | | |
|---|---|---|---|---|
| | a | b | $\mathtt{s}_1$ | $\mathtt{s}_2$ |
| a | 1/16 | 1/16 | 1/16 | 1/16 |
| b | 1/16 | 1/16 | 1/16 | 1/16 |
| $\mathtt{s}_1$ | 1/16 | 1/16 | **1/8** | **0** |
| $\mathtt{s}_2$ | 1/16 | 1/16 | **0** | **1/8** |

Table S4.1: Probability of each pair of tokens in the same sequence.

1. One of two individuals generate a sequence at random (uniformly sample the individual-specific token $s \in \{\mathtt{s}_1, \mathtt{s}_2\}$).
2. Uniformly randomly choose whether a shared or individual-specific token is sampled.
3. Sample one of two shared tokens ($x \in \{\mathtt{a}, \mathtt{b}\}$) or emit the individual-specific token ($= s$).

Then, the probability of each pair of predecessor and follower tokens is as shown in Table S4.1 regardless of their distance; we will encounter every possible pair at random except that only one of the two individual-specific tokens is included in a single sequence, and thus, we will never see the heterogeneous pairs, $(\mathtt{s}_1, \mathtt{s}_2)$ and $(\mathtt{s}_2, \mathtt{s}_1)$. Accordingly, the mutual information, if measured globally across sequences, is constant at 0.25. This contradicts the agent-oriented concept of context dependency where the individual agents generate the sequences without referring to the past tokens, and those who read and/or hear the sequences can predict which one of $\mathtt{s}_1$ and $\mathtt{s}_2$ will come next based on their latest occurrence, not further past.

#### S4.2 Algebraic Complexity of the Corrected Mutual Information Analysis

The particular problem with individual-specific tokens discussed above is not difficult to solve because we can simply condition the relevant probabilities (Eq. 5) on the individual that generate each sequence. However, the mismatch between the mutual information and the agent-oriented concept of context dependency is not limited to that specific case. In this section, we show that data collected from finite-state automata (FSA), which has been a popular model of Bengalese finch song (Hosino and Okanoya, 2000; Okanoya, 2004; Kakishita et al., 2007), can yield different mutual information scores despite their identical context dependency from a generative perspective. The simulations in this section are complex and it was not easy to obtain the mutual information from the formal definition in Eq. 5. Thus, we used the version of mutual information proposed by Sainburg et al. (termed "SMI" below 2019a, cf. Futrell et al. 2019 applied the same metric to hierarchical dependencies in human language syntax) for analysis of real birdsong data, which computes an estimated mutual information $\hat{I}$ of data (Grassberger, 2003; Lin and Tegmark, 2017) and corrects it with shuffled data $X_{\text{sh}}, X_{\text{sh},+d}$.[6]

$$\begin{aligned}
\hat{I}(X, X_{+d}) :=& \hat{S}(X) + \hat{S}(X_{+d}) - \hat{S}(X, X_{+d}) \\
\hat{S} :=& \log_2 N - \frac{1}{N} \sum_x N_x \frac{\psi(N_x)}{\log 2} \\
\text{SMI} :=& \hat{I}(X, X_{+d}) - \hat{I}(X_{\text{sh}}, X_{\text{sh},+d})
\end{aligned}$$

where $N_x$ counts the occurrences of the category or category pair $x$, and $N := \sum_x N_x$. Note that $\hat{S}$ estimates the entropy by approximating $\log N_x$ by $\psi(N_x)$, which is considered robust against data sparsity compared to the naive estimation (Grassberger, 2003). SMI can capture the concept of context dependency in a better manner owing to the utilization of shuffled baseline in its computation. For example, SMI will be zero for the example with individual-specific tokens discussed in the previous section. The mutual information does not change during the shuffling operation, so the score of the original data is canceled out by that of the shuffled data.

[6]The shuffling operates on each sequence; therefore, tokens in different sequences are not mixed.

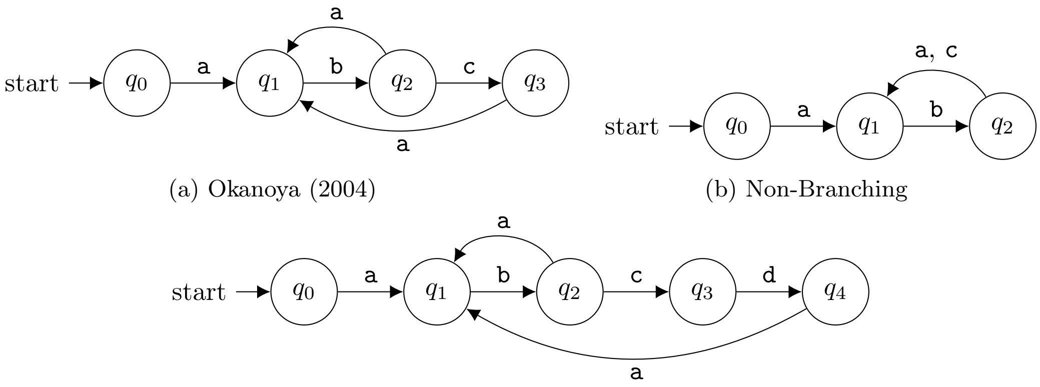


(a) Okanoya (2004)

(b) Non-Branching

(c) Extended

Figure S4.2: FSA models of Bengalese finch song. (a) FSA proposed by Okanoya (2004). (b) A simplified version of (a), removing the transitional branch at $q_2$ while keeping the two possible emissions, `a` and `c`. (c) An extended version of (a), delaying the loop back to $q_1$ after the choice of `c`.

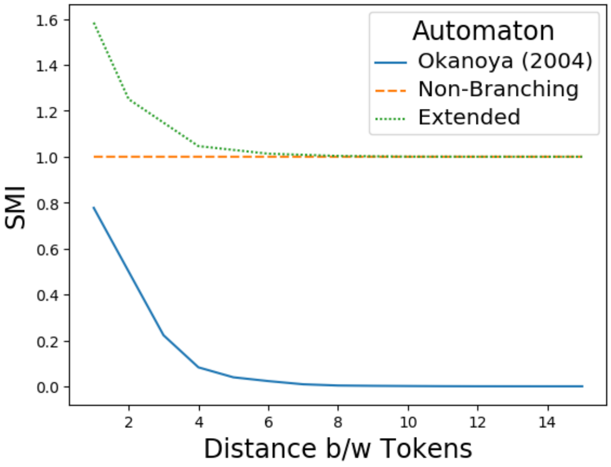


Figure S4.3: SMI of time series data generated by FSA in Figure S4.2.

However, there are other cases where SMI diverges from the agent-based concept of context dependency. In the rest of this section, we discuss *periodic* Markov processes (Lin and Tegmark, 2017). A popular birdsong model (and voice sequences of other animals) is finite-state automata (FSA; Hosino and Okanoya, 2000; Okanoya, 2004; Kakishita et al., 2007, but see Kershenbaum et al., 2014 and Morita and Koda, 2019 for possible effectiveness of language models beyond the capacity of FSA). FSA transitions among a finite number of states and emits/processes a token associated with the transition. Figure S4.2a shows a FSA model of a Bengalese finch song that was proposed by Okanoya (2004) and is commonly cited in studies about the song syntax (e.g., Berwick et al., 2011; Miyagawa et al., 2013). We generated a sequence of $100,000$ tokens from this FSA (with the uniformly random choice between `a` and `c` at the state $q_2$), and the SMI estimated from this data is shown in Figure S4.3. While the SMI dropped exponentially fast, it is hard to recover the Markovian order of the model, or the context dependency from an agent's perspective: it was not until the inter-token distance was 8 or greater that the SMI went below 0.01 while an agent only needs to remember the last emitted token to correctly simulate the FSA (i.e., the FSA can be simulated by a bigram model; speaking in the language of formal language theory, the generated patterns are 2-strictly locally testable).[7]

[7] Berwick et al. (2011) argue that the formal language characterized by the FSA in Figure S4.2a is not strictly locally testable. This is incorrect because we can test whether each string is a possible outcome of the FSA simply by matching each of its substrings of length 2 to the repertoire. {`ab`, `ba`, `bc`, `ca`}.

Moreover, small modifications to the FSA in Figure S4.2a can result in completely different SMI scores. Figure S4.2b removes the branch at $q_2$ and makes the loop back to $q_1$ obligatory, while there are still two possible followers of `b` (= `a` and `c`) chosen at random. While this change does not require any extra effort for the data production/processing, and could even simplify the process owing to the reduced number of states, the SMI is now constant around 1.0. Likewise, the small extension shown in Figure S4.2c, delaying the loop back to $q_1$ after the choice of `c`, puts the SMI convergence around 1.0. This extension does not change the agent-based context dependency, either as agents can still simulate the extended FSA if they remember the last emitted token and nothing further from the past, preserving the 2-strictly local testability. The non-zero SMI of the non-branching and extended FSAs roots in their periodicity.[8] Taking the non-branching FSA (Figure S4.2b) for example, we only observe the predecessor-follower pairs, $(\mathtt{a}, \mathtt{c})$, $(\mathtt{c}, \mathtt{a})$, and $(\mathtt{b}, \mathtt{b})$, when they are $2m$ tokens apart ($m \in \mathbb{Z}_+$), and the other patterns occur elsewhere ($2m+1$). Thus, the joint probability of the pairs is different from the product of each member's marginal probability, pulling $I$ (and $\hat{I}$) above zero. On the other hand, this periodicity is broken by the shuffling, creating a difference between $\hat{I}(X, X_{+d})$ and $\hat{I}(X_{\text{sh}}, X_{\text{sh},+d})$ and keeping the SMI non-zero. Similarly, the extended FSA in Figure S4.2c goes back to each state two to four steps after it leaves the state. Hence, we never see homogeneous pairs like $(\mathtt{b}, \mathtt{b})$ when their distance is an odd integer.

By contrast, the SECL of the three automata was all 1, which matched the Markovian order of the automata without over-estimating the context dependency. While it is not clear how much periodicity exists in real birdsong and other sequential data in biology, potential differences between the mutual information and model-based analysis should be recognized. Thus, we conclude that mutual information cannot replace the model-based analysis for the assessment of agent-oriented context dependency.

### S5 Detailed Comparison with the Markovian Analysis of Context Dependency

Previous studies on context dependency in birdsong often modeled song syntax by a Markov process (Katahira et al., 2011; Markowitz et al., 2013). In this supplementary section, we discuss limitations of this approach, with a specific focus on the method proposed by Markowitz et al. (2013).

#### S5.1 Preliminaries

Markowitz et al. (2013) estimated the length of context dependency in canary song (whose tokens are chunks of syllables, called *phrases*) based on an algorithm that initially assumes an empty context (i.e, tokens are generated independently of previous outputs) and incrementally checks whether or not longer contexts are needed (under an arbitrary upper bound; Ron et al., 1996; Bejerano and Yona, 2001). The algorithm can assign different Markov orders to different contexts; for example, production of a binary signal can be dependent on two previous outputs (second order Markov process) when the most recent one is `0` (i.e., $\mathbb{P}(x \mid \mathtt{00}) \neq \mathbb{P}(x \mid \mathtt{10})$) but not otherwise (i.e., $\mathbb{P}(x \mid \mathtt{01}) = \mathbb{P}(x \mid \mathtt{11})$). The necessity of a longer context is judged based on two types of probabilities: the joint probability of the context, $\mathbb{P}(x_t \mid x_{t-L}, \dots, x_{t-1})$, and the conditional probability of a next token $x_t$ given the context, $\mathbb{P}(x_t \mid x_{t-L}, \dots, x_{t-1})$. The joint probability must be above a threshold $\theta_{\text{joint}}$, filtering out infrequent contexts as negligible. Then, the conditional probability based on the non-infrequent contexts are thresholded ($\mathbb{P}(x_t \mid x_{t-L}, \dots, x_{t-1}) \geq \theta_{\text{trans}}$), and it must also be substantially different from the probability of the same successor token conditioned on the shorter context ($\mathbb{P}(x_t \mid x_{t-L}, \dots, x_{t-1}) / \mathbb{P}(x_t \mid x_{t-L-1}, \dots, x_{t-1}) \geq r \vee \mathbb{P}(x_t \mid x_{t-L}, \dots, x_{t-1}) / \mathbb{P}(x_t \mid x_{t-L-1}, \dots, x_{t-1}) \leq 1/r$). Contexts passing these tests are considered necessary.

Markowitz et al.'s algorithm has two major difficulties in practical applications. The first is the estimation of the joint and conditional probabilities. The number of possible contexts grows exponentially as they get longer, and fewer observations of each become available. Thus, we cannot naively estimate a distinct probability distribution per long context due to the data shortage and some generalizations over different contexts are needed. In classic Markovian modeling, contexts were often generalized according to their suffix substrings so that the full order conditional probability was estimated from lower order statistics (Katz, 1987; Kneser and Ney, 1995; Goldwater et al., 2006; Teh, 2006). Markowitz et al.'s algorithm is also one such example wherein unnecessarily long contexts are reduced to their suffixes. Even using such advanced

[8] Lin and Tegmark (2017) prove that mutual information in periodic Markovian processes is characterized by an exponential decay plus a constant bottomline.

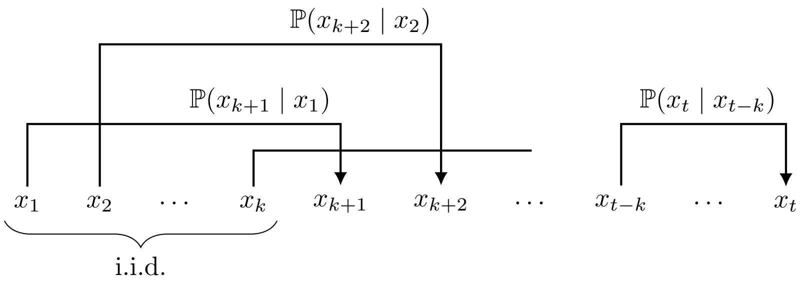


Figure S5.1: Schematic diagram of the Markov process with $k$-step delays.

techniques, however, Markovian models were only able to scale up to a few order; for example, even Google's language model—exploiting their big corpora—was based on five-grams (i.e., fourth order Markov; Michel et al., 2011). Similarly, Markowitz et al. estimated the dependency length in canary song as seven, but this was the upper bound set for the algorithm to run. By contrast, recent models based on artificial neural networks represent discrete contexts in a continuous-valued space, wherein generalizations across contexts are made more flexibly (Bengio et al., 2001, 2003). This innovation made it possible to process long context dependencies that potentially range over hundreds of tokens (Khandelwal et al., 2018; Dai et al., 2019).

The second problem with Markowitz et al.'s method is the difficulty in tuning hyperparameters. As introduced above, the algorithm is parameterized by three thresholds $(\theta_{\text{joint}}, \theta_{\text{trans}}, r)$ as well as the upper bound $L_{\text{MAX}}$ on the possible Markov orders. As we will demonstrate in the next section, different settings of these parameters lead to completely different results and no lesson about their optimization is provided in the literature. Here, our Transformer-based estimation of context dependency has an advantage of robustness; in the next section, we will recover the correct dependency length behind simulated data *using exactly the same hyperparameters as in our birdsong analysis.*

#### S5.2 Comparison Using Simulated Data

This section demonstrates the problems with the Markovian estimation of context dependency and advantages of our proposed method based on Transformer language modeling, using simulated data. The data are generated by a delayed Markov process, which is schematized in Figure S5.1; each token is sampled conditioned on the $k$-th most recent token in the context (and nothing else); initial $k$ tokens are i.i.d. We adopted the uniform distribution over 37 symbols (= the number of Bengalese finch syllable categories estimated by the ABCD-VAE) for the initial i.i.d. sampling. The transitional probabilities was uniform over three symbols that were randomly selected from the 37 symbols for each condition at the beginning and fixed throughout the sampling process. We set $k = 2, 4, 8, 16$, and for each $k$, we sampled 10,000 sequences as training data, each of which consisting of 128 tokens, and 100 sequences of the same length as test data (only used in our Transformer-based method).

The maximum Markovian order for Markowitz et al.'s method was set as $L_{\text{MAX}} = 20 > k$. For the other three parameters, we examined two sets of values used in the literature:

- Markowitz et al. (2013)
  - $\theta_{\text{joint}} = 0.007$
  - $\theta_{\text{trans}} = (1 + 17.5) \times 0.01 = 0.185$
  - $r = 1.6$
- Bejerano and Yona (2001)
  - $\theta_{\text{joint}} = 0.0001$
  - $\theta_{\text{trans}} = (1 + 0) \times 0.001 = 0.001$
  - $r = 1.05$

Table S5.1: Maximum length of effective contexts detected by the Markovian method. The algorithm was run using two different sets of parameter values, one adopted by Markowitz et al. (2013) and the other by Bejerano and Yona (2001).

| Delay ($k$) | Maximum Context Length Detected | |
|---|---|---|
| | Markowitz et al. (2013) | Bejerano and Yona (2001) |
| 2 | 0 | 5 |
| 4 | 0 | 3 |
| 8 | 0 | 3 |
| 16 | 0 | 3 |

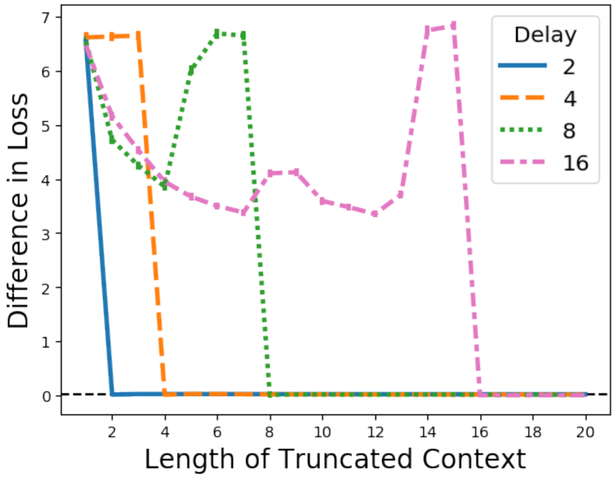


Figure S5.2: The differences in the mean loss (negative log probability) between the truncated- and full-context predictions of time series generated by a $k$-step delayed Markov process ($k = 1, 4, 8, 16$). The x-axis corresponds to the length of the truncated context. The error bars show the 90% confidence intervals estimated from 10,000 bootstrapped samples. The loss difference is statistically significant if the lower side of the intervals are above the threshold indicated by the horizontal dashed line.

The maximum length of contexts that were judged as necessary under each of the two settings is reported in Table S5.1. When we adopted Markowitz et al.'s parameters, no effective context was detected. Their threshold for the joint probability, $\theta_{\text{joint}}$, was so large that only singleton contexts were qualified. For the same reason, context dependency was underestimated under Bejerano and Yona's settings for $k = 4, 8, 16$; long contexts were not probable enough to pass the threshold. Note that naively decreasing $\theta_{\text{joint}}$, and thereby being more tolerant for infrequent conditions, does not lead to the true dependency. This was evidenced by the $k = 2$ example, whose dependency length was overestimated under Bejerano and Yona's settings. Given these results, we conclude that the Markovian estimation of context dependency is not robust.

In contrast to the Markovian estimation, our proposed method based on Transformer language modeling correctly recovered the true dependency length of the delayed Markov process for $\forall k = 2, 4, 8, 16$ (Figure S5.2). The difference between the truncated- and full-context predictions of the test data by the Transformer language model vanished when the length of the truncated context matched $k$ (i.e., the SECL was equal to $k$). Note that we used the same hyperparameters across different datasets, including birdsong, suggesting that the proposed method is more robust than the previous Markovian estimation.
